# Salinity impacted microbial carbonate precipitation mechanisms of modern microbialites in peritidal zone

**DOI:** 10.1101/2024.11.11.623098

**Authors:** Yunli Eric Hsieh, Sung-Yin Yang, Shao-Lun Liu, Shih-Wei Wang, Wei-Lung Wang, Sen-Lin Tang, Shan-Hua Yang

**Author notes:** corresponding author: Shan-Hua Yang.

## Abstract

Microbialites, ancient records of microbial activity, serve as significant indicators of environmental change. This study examined microbialites from the peritidal zone of three tide pools at Fongchueisha, Hengchun, Taiwan, to investigate the impact of salinity on microbial community composition and carbonate precipitation mechanisms. Microbial samples were collected across varying salinity gradients over multiple timepoints and analyzed using next-generation sequencing of bacterial 16S and eukaryotic 18S rRNA genes. Our findings reveal that the microbial communities in higher salinity environments exhibited significant shifts, with increased relative abundance of ureolytic bacteria and ammonifying microorganisms, such as *Myxococcota* and *Actinobacteriota*. This suggests the presence of diverse microbial carbonate precipitation mechanisms beyond photosynthesis, including ureolysis and ammonification. Furthermore, our results show that changes in the composition of cyanobacteria and diatoms were influenced by salinity, with heterocystous cyanobacteria (e.g., *Nostocales*) dominating low-salinity environments, and non-heterocystous cyanobacteria (e.g., *Synechococcales*) prevailing in higher salinity environments. Functional predictions reveal that microbial communities in high-salinity environments were enriched in anaerobic metabolic pathways, including pyruvate fermentation and the urea cycle. These findings highlight the significant role of salinity in influencing microbial composition and metabolic pathways, shaping carbonate precipitation processes in microbialites.

**Importance:** The study focuses on the impact of environmental salinity on microbial community composition and carbonate precipitation mechanisms within modern microbialites, based on an analysis of samples from three tide pools with different salinity levels collected at five time points throughout the year. Using next-generation sequencing of bacterial 16S and eukaryotic 18S rRNA genes, we identified key shifts in microbial communities along salinity gradients and explored diverse microbial processes involved in carbonate precipitation. This work enhances our understanding of microbial ecosystems within modern microbialites and their response to environmental changes. Additionally, our study provides insight into ancient biogeochemical processes, with implications for interpreting microbial metabolism in carbonate precipitation across different salinity regimes.

## Introduction

Lithified deposits formed by microbial activity (microbialites) represent the most continuous and ancient record of life on Earth, having persisted as fossils for at least 3.45 billion years (1, 2). Based our limited understanding, modern microbialites are confined to areas where competitors and destructors are absent or where the biogeochemical conditions are favorable for their consistent growth (3). Although many studies on modern microbialites in coastal and inland area have been conducted in recent years, microbialites that form exclusively in the supratidal zone under fresh to normal seawater conditions represent a distinct type that has received little attention (4–6). The formations of such microbialites are characterized by their proximity to the ocean: microbialites in the upper formations receive freshwater from inflow seeps, those in the middle formations endure a mix of freshwater seepage and marine overtopping, and the lower formations are in closest contact with the ocean (7). G. M.

Rishworth et al. (3) proposed that the presence of these microbialites across three continents and their recent discovery suggest they may be more common than previously thought, as they might have been overlooked along other coastlines.

Most records of modern microbialites in the peritidal zones are found in South Africa, Western Australia, and Northern Ireland (3). However, modern microbialites have been observed in the peritidal zone of the Fongchueisha area in Hengchun, located in southern Taiwan. This region, characterized by a tropical monsoon climate, a long and varied coastline, and diverse ecological habitats—including limestone terrain and coral reefs—appears to be the only location in Taiwan where such microbialite structures have been reported. Notably, modern microbialites in Fongchueisha’s peritidal zone thrive near freshwater sources closer to land, but as one moves toward the sea, their development diminishes. This observation implies that the ratio of seawater to freshwater could be a key factor in limiting microbialite growth.

Furthermore, from a microscopic perspective, the formation of microbialites, T. Zhu and M. Dittrich (8) mentioned that the proportion of microbes causing carbonate precipitation varies in different environments. Compared to soil, caves, hypersaline lakes, and hot springs, freshwater and marine systems have the most diverse species and mechanisms of microbial carbonate precipitation, including photosynthesis, ureolysis, denitrification, sulfate reduction, and anaerobic oxidation of methane. In freshwater systems, *Cyanobacteria* and other photosynthetic microbes dominate, whereas in marine systems, photosynthesis, sulfate reduction, and anaerobic oxidation of methane are more prominent (8). Besides, it has been found that bacteria may use ureolysis for microbial carbonate precipitation in high saline environments (9). However, given that modern microbialite in the peritidal zone are influenced by both freshwater and seawater, it raises the question of whether other microbial carbonate precipitation mechanisms and microbes have been overlooked in favor of photosynthetic microbes, that have historically received more attention.

In this study, we collected microbialite samples from three tide pools in the Fongchueisha area of Hengchun, spanning from the freshwater-influenced supralittoral zone to the seawater-influenced lower intertidal zone, over the course of a year. We analyzed the V9 region of the 18S rRNA gene for eukaryotes and the V6-V8 region of the 16S rRNA genes for bacteria using next-generation sequencing to understand the temporal and spatial variations in the composition of lithic microbes. Through this, we aim to determine the main possible mechanisms of microbial lithification in intertidal stromatolites influenced by different proportions of seawater and freshwater.

## Results

### Morphological variations of microbialites in tide pools

The scanning electron microscope (SEM) analysis revealed distinct morphological variations among the microbialites from the three examined tide pools (Fig. S1). Moreover, the morphological differences between the surface and bottom microbialites within each tide pool were also observed, with the most pronounced disparity occurring in tide pool 30. Specifically, while the microbialite structures on the surface and bottom of tide pools 0, 12, and the surface of tide pool 30 exhibited more complex architectures, the microbialite structure at the bottom of tide pool 30 was notably simpler. These observations suggest that the microbialite formations at the bottom of tide pool 30 are distinct from those found in other locations.

### Influence of environmental factors on bacterial alpha diversity and community composition in tide pools

The correlation between salinity and bacterial alpha diversity in microbialites from the three tide pools indicates that the disparity between the surface and bottom of tide pool 30 is more pronounced than in the other two tide pools (Fig. 2). Specifically, the salinity at the bottom of tide pool 30 is the highest, while it is comparatively lower in tide pools 0 and 12 (Fig. 2a, c). At the surface, the salinity of tide pool 30 is similar to that of tide pools 0 and 12 (Fig. 2b, d). However, there is no significant correlation between bacterial alpha diversity indices and salinity across the tide pools. These findings suggest that, while salinity differences highlight the distinct nature of the bottom environment in tide pool 30, bacterial alpha diversity does not reflect these differences.

**Figure 1.**
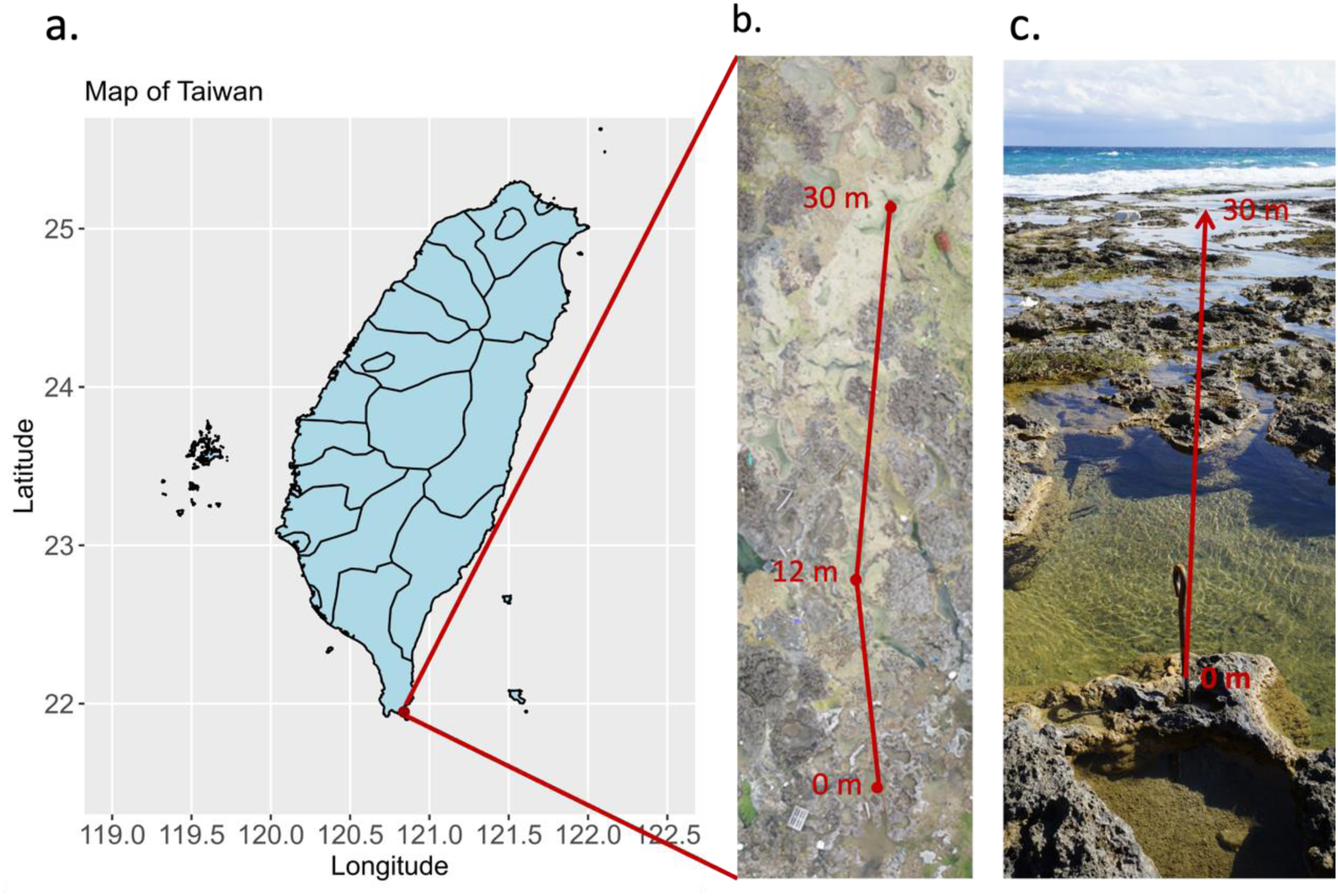
Sampling sites. (a) the map of Taiwan and the sampling site is located in Hengchun. (b) Aerial photography shows that the distance among three tide pools in the peritidal zone. (c) the distance between the tide pool 0 and tide pool 30.

**Figure 2.**
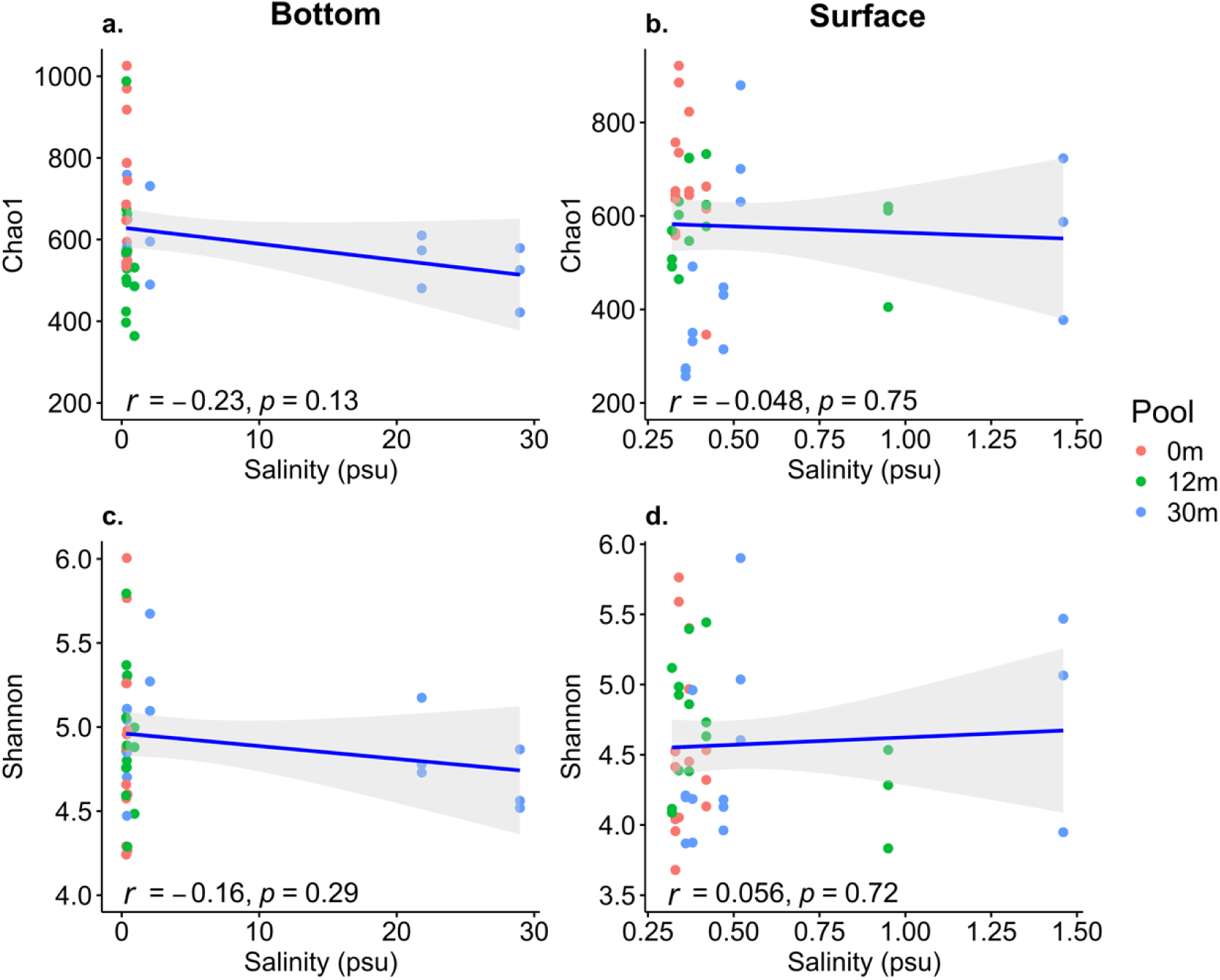
Correlation between salinity and bacterial alpha diversity of peritidal zone modern microbialites in three tide pools. (a) and (b) are the correlation between salinity and bacterial Chao1 diversity index of bottom and surface parts respectively. (c) and (d) are the correlation between salinity and bacterial Shannon diversity index of bottom and surface parts respectively.

Regarding temperature, there was minimal variation between the surface and bottom across the three tide pools (Fig. S2a-d). Dissolved oxygen levels were higher at both the surface and bottom of tide pool 30 compared to the other two tide pools (Fig. S2e-h). In terms of Total Dissolved Solids (TDS) and Suspended Particulate Concentration (SPC), tide pool 30 exhibited significant differences between its surface and bottom, particularly at the bottom, compared to the other tide pools (Fig. S2i-p). The pH levels were higher in tide pool 30, while tide pool 12 demonstrated the greatest variability (Fig. S2q-t). These results, combined with the earlier salinity findings (Fig. 2), indicate that tide pool 30 shows the most pronounced differences between its surface and bottom environments, with the bottom characterized by higher salinity, TDS, and SPC.

In analyzing the correlation between bacterial alpha diversity and environmental factors, no significant correlation was found between the four environmental factors at the surface of the tide pools and bacterial alpha diversity. However, at the bottom of the tide pools, a significant negative correlation was observed between temperature and the Shannon diversity index (*r* = -0.38, *p* < 0.05; Fig. S2c). Additionally, dissolved oxygen displayed a significant negative correlation with bacterial Chao1 (R = -0.43, *p* < 0.05; Fig. S2e). Canonical correlation analysis (CCA) comparing the relationships between bacterial communities and environmental factors in the tide pools over time (Fig. S3) revealed that the bacterial communities in tide pool 30 showed a more significant correlation with environmental factors than those in the other tide pools. Specifically, in the January and October bottom samples of tide pool 30, bacterial composition was positively correlated with salinity, TDS, PSU, and DO. Overall, although environmental factors exert minimal influence on bacterial alpha diversity, the CCA results suggest that microbial community composition is significantly influenced by environmental factors.

### Eukaryotic and prokaryotic composition variations in tide pools

The analysis of eukaryotic composition in modern microbialites from the peritidal zone across three tide pools (Fig. S4) indicated that Bacillariophyta were the predominant group within Ochrophyta, regardless of their proximity to seawater. Within Bacillariophyta (Fig. S4a), the class Bacillariophyceae, particularly the orders Naviculales and Thalassiophysales, was identified as the dominant group. An increased relative abundance of Naviculales was observed in tide pool 30. Additionally, the class Conscinodiscophyceae showed increased abundance in environments with higher seawater concentrations. In the Chlorophyta group (Fig. S4b), Ulvophyceae were more dominant in areas closer to seawater, while Chlorodendrophyceae and Trebouxiophyceae were detected only in environments with higher freshwater content.

Regarding prokaryotic communities, the main bacterial composition of modern microbialites in the peritidal zone was consistent at the phylum level, with *Cyanobacteria*, *Chloroflexi*, and *Proteobacteria* being the most abundant (Fig. 3a). Notably, *Myxococcota* appeared at the bottom of tide pools 12 and 30, while *Actinobacteriota* were more prevalent at the bottom of tide pool 30. At the class level, the differences in bacterial composition between the surface and bottom layers were more pronounced in areas closer to the seaside. Specifically, within the surface bacterial community of tide pool 30, *Chloroflexia* exhibited a higher relative abundance compared to the surface layers of tide pools 0 and 12 (Fig. 3b-d). These findings suggest that environmental factors, particularly salinity, TDS, PSU, and DO, have more influence on *Myxococcota*, *Actinobacteriota*, and the class *Chloroflexia* within the *Chloroflexi* phylum.

**Figure 3.**
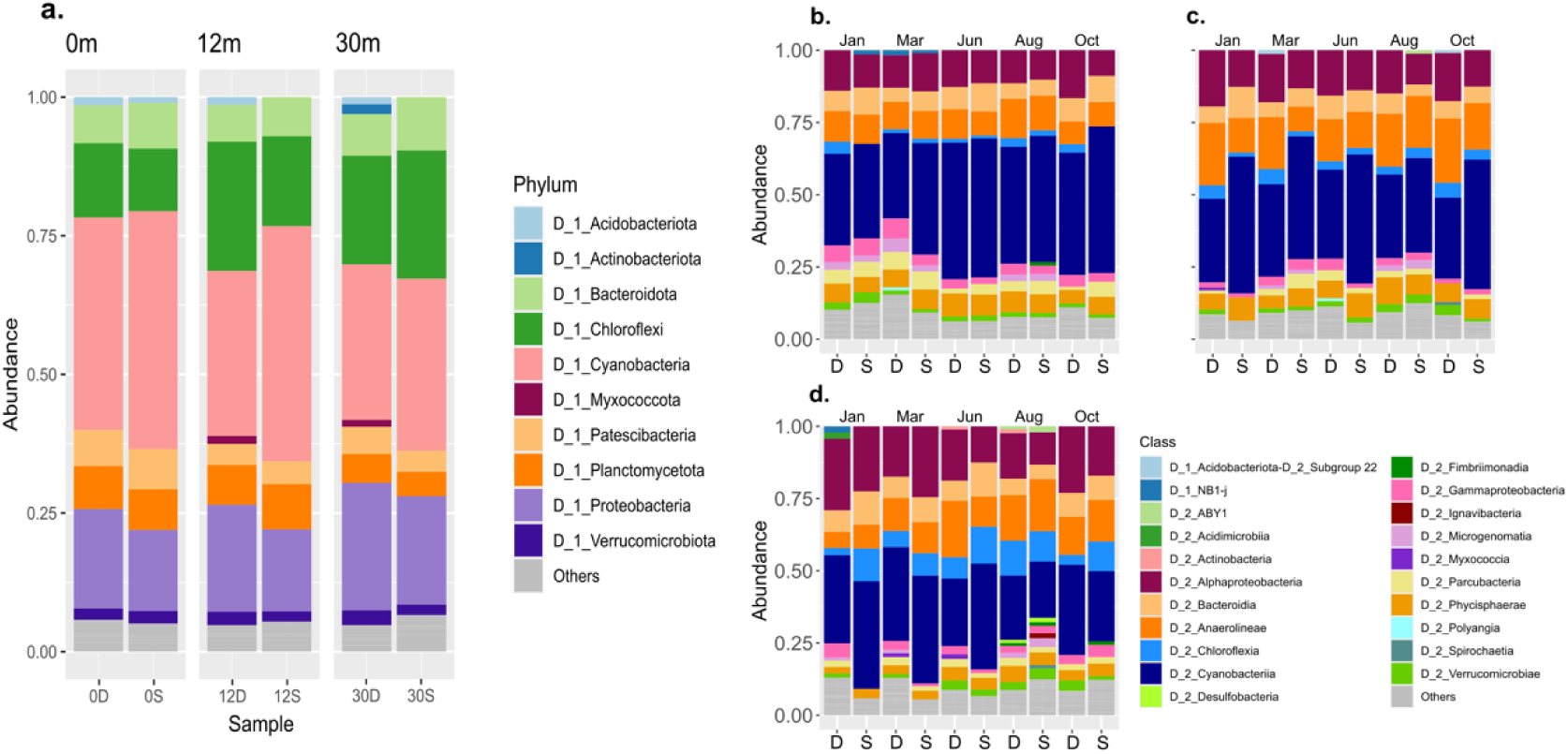
Bacterial compositions in peritidal zone modern microbialites of tide pools. (a) Bacterial compositions of phylum level. (b-d) Bacterial compositions in class level during five sampling time. (b) tide pool 0; (c) tide pool 12; (d) tide pool 30. S indicates surface part of the pools, and D indicates bottom part of the pools.

Within the *Proteobacteria* phylum (synonym *Pseudomonadota*), *Alphaproteobacteria* was identified as the most dominant class. The composition of different families within *Alphaproteobacteria* varied according to the specific tide pool and the sampling time (Fig. 4). In tide pool 30, where salinity levels were the highest, the relative abundance of *Rhodobacterales* increased. Conversely, *Rhizobiales* exhibited the highest relative abundance in tide pool 12. In tide pool 0, *Sphingomonadales* displayed a higher relative abundance compared to the other tide pools.

**Figure 4.**
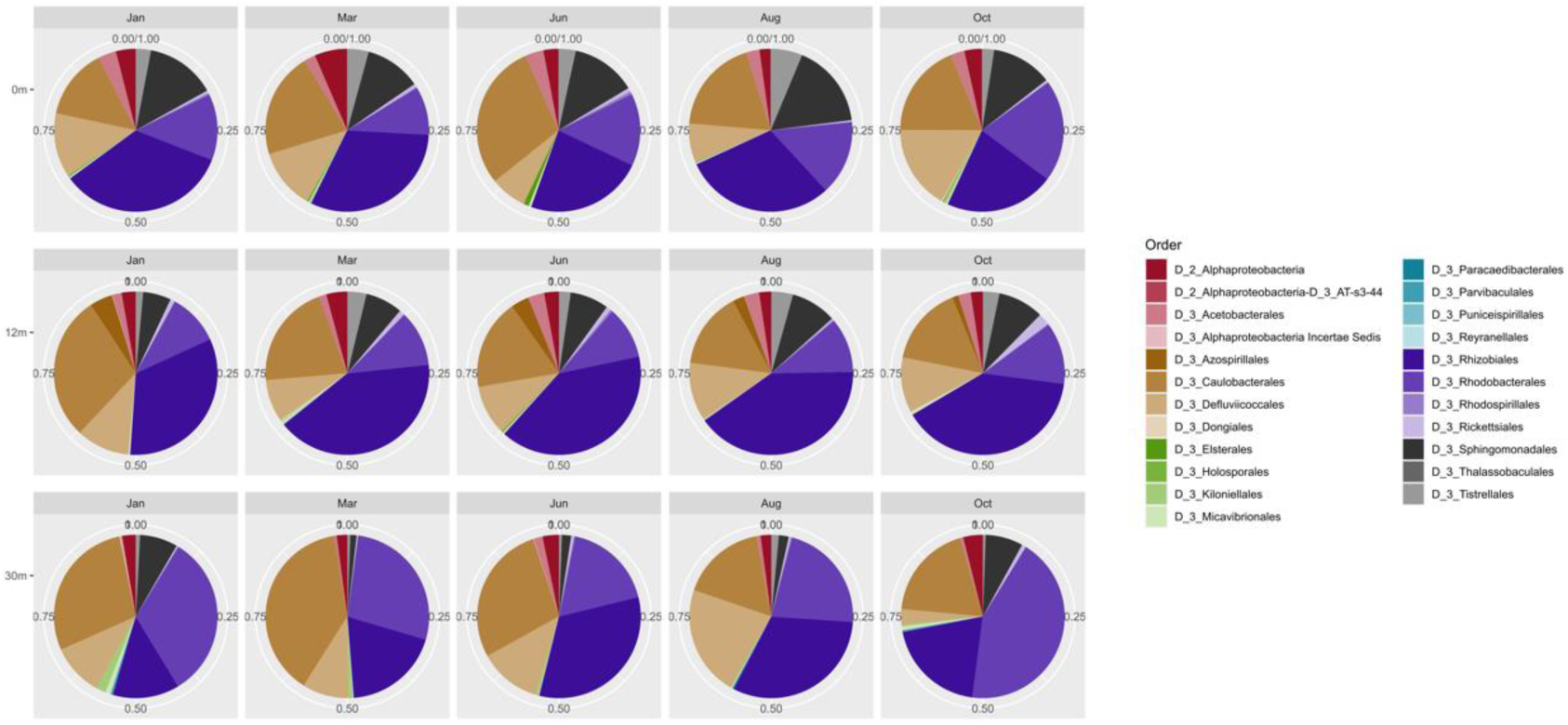
The changes in the populations within the *Alphaproteobacteria* of modern microbialites in different tide pools across five different months. Colors indicates different orders in the class.

The composition of different families within *Cyanobacteria* also varied depending on the tide pool. At different time points in tide pool 0, the family *Nostocaceae*, a group of heterocyst-forming *Cyanobacteria*, was predominantly dominant. In tide pool 12, both *Nostocaceae* and *Oxyphotobacteria* incertae sedis were dominant groups, with their relative abundances fluctuating seasonally (Fig. 5b). Phylogenetically, *Oxyphotobacteria* incertae sedis is closer to the filamentous families within *Synechococcales*, suggesting that this family consists of filamentous *Cyanobacteria* without heterocysts (Fig. 5a). In tide pool 30, *Oxyphotobacteria* incertae sedis generally exhibited higher relative abundance than *Nostocaceae*. These findings indicate that in the peritidal zone, the dominant *Cyanobacteria* are primarily filamentous, and proximity to seawater is associated with a decrease in the relative abundance of heterocyst-containing *Cyanobacteria*.

**Figure 5.**
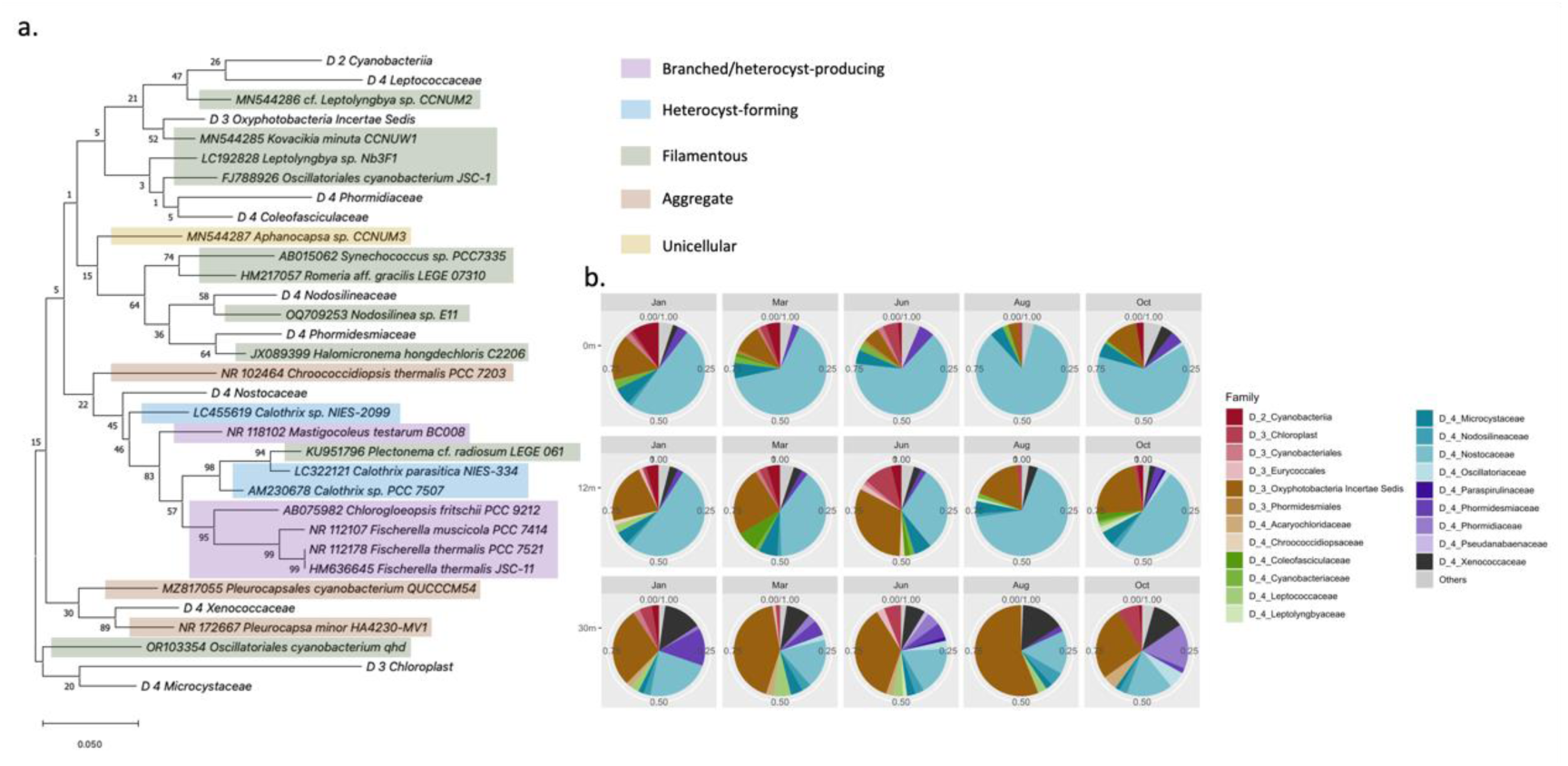
Phylogenetic tree and dynamics of cyanobacterial community in each in the peritidal zone modern microbialites during five sampling time. In phylogenetic tree, the different morphologies of cyanobacteria are labelled with different colors. (b) The changes in the populations within the cyanobacteria of modern microbialites in different tide pools across five different months.

Within the *Nostocaceae* family, there were clear differences in composition between the surface and bottom layers, particularly in tide pool 12 (Fig. S5), where *Rivularia* predominantly composed the bottom layer. *Rivularia* was also the main genus of *Nostocaceae* in tide pool 30. Compared to the other two tide pools, tide pool 12 exhibited more pronounced seasonal differences in the composition of the *Nostocaceae* family, resembling tide pool 30 more in winter and tide pool 0 more in summer (Fig. S5). This finding suggests that, despite the *Cyanobacteria* group being the most dominant in each tide pool, the composition of these communities is subject to change with environmental fluctuations.

### Bacterial network associations with dominant Cyanobacterial genera

The bacterial network analysis revealed associations between bacterial genera and the dominant cyanobacterial genera based on their relative abundance. In the bacterial network of tide pool 0 (Fig. S6a), *Nostocaceae* exhibited numerous connections with other bacteria, whereas *Oxyphotobacteria* incertae sedis showed fewer connections. Conversely, in the network of tide pool 30 (Fig. S6b), *Nostocaceae* had fewer connections with other bacteria, while *Oxyphotobacteria* incertae sedis displayed more extensive connections. These findings suggest that the bacterial composition within the microbial mats of microbialites may be influenced by the relative abundance of dominant cyanobacterial genera. Furthermore, the previously identified *Chloroflexia* (*Chloroflexi*) and *Myxococcaceae* (*Myxococota*) were also associated with the dominant *Cyanobacteria*, indicating potential interactions within these microbial communities.

### Functional predictions of bacterial communities across tidal pools

To investigate functional changes in the bacterial communities, we utilized PICRUSt2 to predict the potential functions of bacteria from each tidal pool. The principal component analysis (PCA) revealed a clear functional shift from the land side (0m) to the seaside (30m) (Fig. 6). Among the top 50 potential functions with significant differences between tidal pool 0 and 30 (Fig. S7), the pathways of pyruvate fermentation to propanoate I and acetyl-CoA fermentation to butanoate II were predominantly enriched at the tidal pool 30. In contrast, the benzoyl-CoA degradation I (aerobic) pathway was mainly enriched at the tidal pool 0. This suggests that bacteria inhabiting tide pool 30 are likely to rely more on anaerobic metabolic pathways. Additionally, the pathways related to the urea cycle was also predominantly enriched in tide pool 30. This enrichment may be associated with microbial carbonate precipitation (MCP), potentially linked to microbial metabolic processes of ureolysis (10).

**Figure 6.**
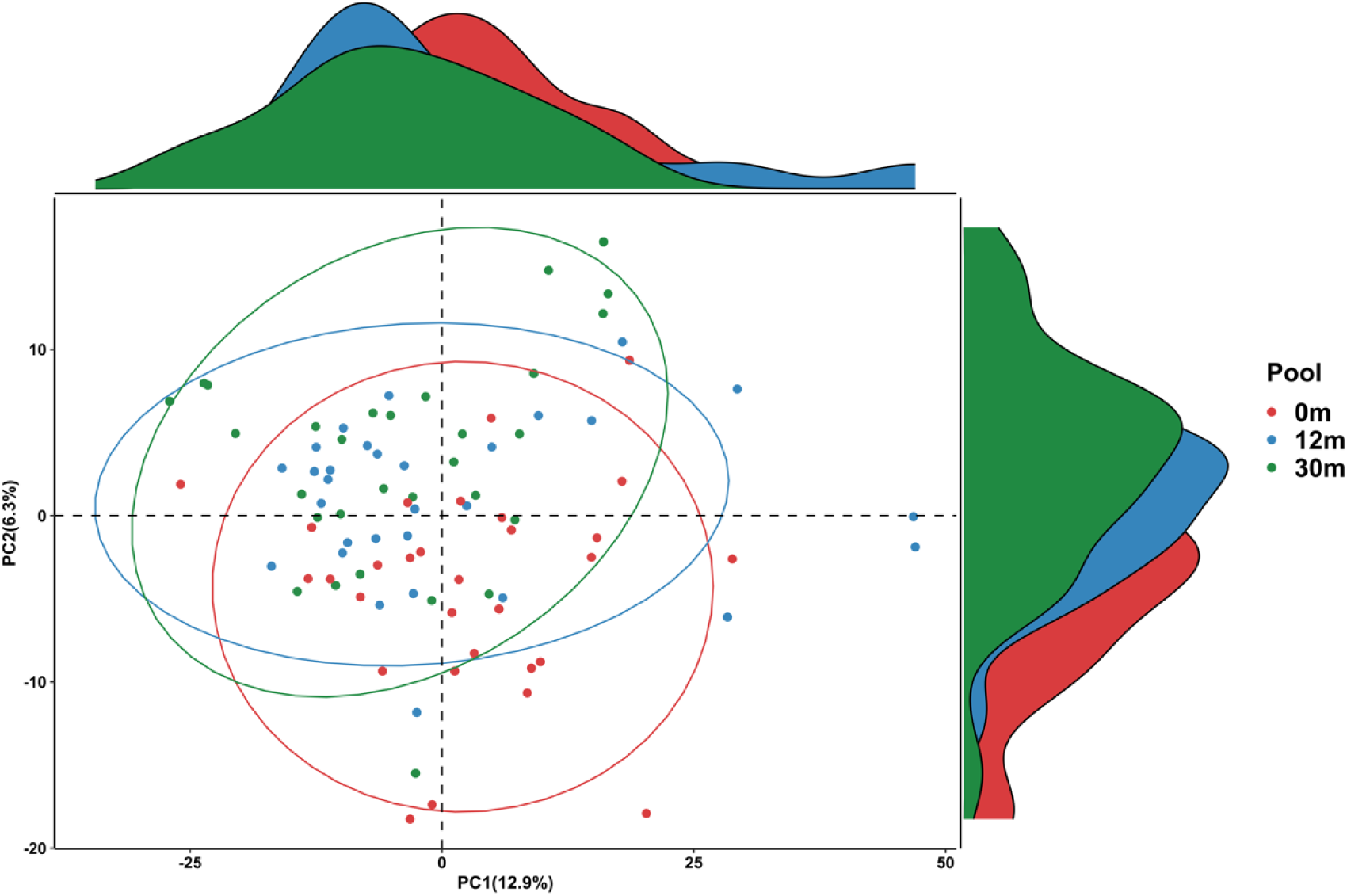
Principal component analysis (PCA) of the potential functions of bacteria in modern microbialites from the peritidal zone across tide pools. Different colors represent different tidal pools: red (0m), blue (12m), and green (30m). The difference between groups was tested by using PERMANOVA (*p* = 0.016).

## Discussion

Based on our observations, we preliminarily infer that microbial involvement in early diagenesis in the Fongchueisha intertidal zone requires the following natural conditions: exposure to the Pacific Ocean and the influence of continuous waves from the open sea, along with relatively rapid tectonic uplift, which creates a wider intertidal zone. Coastal currents and the northeast monsoon form sand dunes near cliffs, providing a steady supply of fine sediments. The porosity difference between the Hengchun limestone and the underlying sandstone allows groundwater to seep through geological interfaces. Dunes and vegetation near limestone cliffs ensure a continuous flow of supersaturated carbonate freshwater from springs. This uplifted intertidal zone, featuring biogenic reefs and bioclastic limestone, is subject to marine, rain, and spring waters, leading to extensive cementation and diagenesis, with spring water lingering in tide pools. Despite these areas being well above mean sea level, the combined effects of Pacific waves and tides create brackish water environments.

### Salinity may affect the composition and changes of microorganisms in peritidal zone modern microbialites, as well as microbial carbonate precipitation mechanisms

It has been known that microbial carbonate precipitation (MCP) can occur as a result of various microbial metabolic processes, including photosynthesis (11), ammonification (12), ureolysis (10), denitrification (13), sulfate reduction (14), and anaerobic oxidation of methane (15). Previous studies on the mechanisms of carbonate precipitation in microbialites have often focused on photosynthesis and photosynthetic microorganisms, such as *Cyanobacteria* (3, 16, 17). However, other mechanisms related to microbial carbonate precipitation, such as ureolysis, denitrification, sulfate reduction, and anaerobic oxidation of methane, have rarely been discussed (9). For instance, in the study by T. Zhu and M. Dittrich (8), it was mentioned that *Myxobacteria* (bacteria within *Myxococcota*) participate in microbial diagenesis through the process of ammonification (the conversion of organic nitrogen to NH_3_). In our results, the relative abundance of *Myxococcota* increases in the bottoms of tide pools 12 and 30.

Besides ammonification, microorganisms can participate in carbonate precipitation through ureolysis. This process involves bacterial urease hydrolyzing urea in the environment into CO_2_ and ammonia, with ammonia dissolving in water and dissociating into ammonium ions. This indirectly raises the microenvironment’s pH, promoting the precipitation of calcium ions as calcium carbonate. In the study by S. T. T. Nguyen et al. (9) on microbialite-forming mats from South Australian saline lakes, the potential for microorganisms to participate in carbonate precipitation through ureolysis was proposed. Our results also show an increase in the relative abundance of *Actinobacteriota* at the bottoms of tide pool 30; some members of *Actinobacteriota* can participate in carbonate precipitation through ureolysis (8).

Additionally, at the tide pool 30, the relative abundance of *Rhodobacteraceae* (*Alphaproteobacteria*) and *Xenococcaceae* (*Cyanobacteria*) was higher. In S. T. T. Nguyen et al. (9)’s study, genes related to ureolysis were enriched in the microbialite-forming mats, primarily coming from bacteria including *Rhodobacteraceae* and *Xenococcaceae*. Additionally, we also found that the pathway of urea cycle was only enriched in the tidal pool 30. Therefore, this study not only finds the presence of ammonification and ureolysis carbonate precipitation mechanisms in peritidal zone modern microbialites but also suggests, through a comparison of tide pools with different salinities, that these two types of carbonate precipitation mechanisms are likely to occur in high-salinity environments.

Additionally, in the peritidal zone modern microbialites, diatoms are common eukaryotic microorganisms (3), and our results also support this phenomenon. Since diatoms possess a urea cycle (18), although it is uncertain whether the CO_2_ and ammonia produced from urea decomposition within their cells directly dissolve in water, we do not rule out the possibility that diatoms may participate in this process to some extent. In future studies on microbial carbonate precipitation in modern microbialites, it is recommended to consider the role of diatoms as a subject of research.

### In peritidal zone modern microbialites, changes in the composition of *Cyanobacteria* may be influenced by salinity

In our results, tide pool 0, which has lower salinity, shows a higher relative abundance of *Nostocales*, while tide pool 30, which is closer to seawater salinity, is dominated by *Synechococcales*. *Nostocles* are cyanobacteria with heterocysts, while *Synechococcales* do not have heterocysts. A. Oren (19) suggested that salinity is unfavorable for heterocytous cyanobacteria in normal marine and hypersaline environments. In addition, the study by M. A. Campbell et al. (17) in Australia also indicated, low salinity favored the presence of heterocytous cyanobacteria in freshwater mats, whereas higher salinities primarily supported the growth of unicellular and filamentous nonheterocytous genera. This aligns with our observations.

M. A. Campbell et al. (17) concluded that environmental factors, rather than the type of mat or time of sampling, influenced the distribution of heterocytous *Cyanobacteria*. They also hypothesized that salinity could be controlling heterocyte glycolipid synthesis in *Cyanobacteria* in intertidal/subtidal zones. Our observations support this hypothesis.

Interestingly, the influence of salinity on community composition does not only occur at higher phylogenetic levels; even within the family *Nostocaceae*, the genera show different compositions based on the salinity of the tide pool. For example, *Rivularia* was the main *Nostocaceae* genus in tide pools with higher salinity. In S. Shalygin et al. (20)’s study on the stromatolites of hypersaline lakes, *Rivularia* was found to be the dominant genus, indicating that *Rivularia* is likely the more salt-tolerant genus within the *Nostocaceae*.

However, our results do not exclude the influence of seasons or time on the microbial composition, including *Cyanobacteria*. Changes in the relative abundance of *Cyanobacteria* and other microorganisms in the three tide pools can still be observed in different months. For example, when the northeast monsoon intensifies, the wind strength affects the amount of seawater entering the tide pools, leading to changes in the salinity of the tide pools. This could subsequently influence the microbial composition in the tide pools, making the microbial composition more similar to that of tide pools exposed to seawater. Such results are especially evident in the transitional tide pool (tide pool 12).

Previous studies have found that the diatoms *Hemiaulus*, *Rhizosolenia*, and *Chaetoceros* form symbiotic relationships with heterocystous cyanobacteria, such as *Richelia* and *Calothrix* (21, 22). These three genera of diatoms belong to the class Bacillariophyceae. In our results, diatoms mainly belong to the class Bacillariophyceae, with a relatively higher abundance in the less saline tide pool 0. Interestingly, in the cyanobacterial composition of this study, *Calothrix* is also dominant in tide pool 0. Although it is currently unclear whether there is a direct relationship between diatoms and *Cyanobacteria* in peritidal zone modern microbialites, this study does not exclude the possibility that nitrogen utilization may be involved in the composition of diatoms and nitrogen-fixing *Cyanobacteria*.

## Conclusion

Microbialites are among the oldest known ecosystems on Earth, with a fossil record extending back over 3.5 billion years. These enduring communities create sedimentary structures through the interaction of microbial carbonate precipitation metabolisms and surrounding environment. Although microbialites were widespread on ancient Earth, modern microbialites are now primarily found in restricted habitats. Studying modern microbialites provides insight into how these ancient ecosystems interact with and respond to environmental changes. It has been known that microbialites serve as indicators of an ever-changing Earth, increasingly affected by global climate change. Therefore, research on modern microbialites offers a unique chance to understand the feedback mechanisms between forming microbialite and changing environment.

In this study, by comparing the bacterial and eukaryotic communities of peritidal zone modern microbialites in environments with different salinities, we believe that in higher salinity environments, in addition to photosynthesis, microorganisms may employ more diverse microbial carbonate precipitation mechanisms, such as ammonification and ureolysis. Furthermore, changes in the composition of *Cyanobacteria* and diatoms may be influenced by salinity, and we suggest that nitrogen utilization may be involved in their composition.

## Materials and Methods

### Microbialite sample and environmental data collection

The microbialite sample were collected at Fongchueisha, Hengchun, Taiwan (24°94’51.1”N, 120°84’25.8”E; Fig. 1). Sampling was conducted along the intertidal zone from land to sea at three tide pools located in the upper (tide pool 0), middle (tide pool 12), and lower (tide pool 30) intertidal zones, with collections made at five time points: June, August, and October 2019, and January and March 2020. For each tide pool, microbialites were sampled from both the surface (S) and bottom (D) of the pools. Following collection, some samples were preserved at 4°C and transported to the laboratory, where they were stored at -80°C for DNA extraction. Additional samples were preserved in a fixative solution (2.5% glutaraldehyde, 4% paraformaldehyde, 0.1 M phosphate buffer) and stored in a dark, cool environment for subsequent electron microscopy analysis.

For environmental parameters, a YSI Pro Plus portable multiparameter water quality instrument was used to measure conditions along the transect. Measurements were taken approximately every 3 meters, from the freshwater source at the upper intertidal zone (tide pool 0) to the lower intertidal zone (tide pool 30). The parameters measured were temperature, salinity (PSU), total dissolved solids (TDS), suspended particulate concentration (SPC), dissolved oxygen (DO), and pH.

### Scanning electron microscope

Sections from the interior of microbialite samples were carefully excised using sterilized blades, and their ultrastructure was analyzed using scanning electron microscopy (SEM). The samples were initially fixed in a solution containing 2.5% glutaraldehyde and 4% paraformaldehyde in 0.1 M phosphate buffer at room temperature for two days. Following fixation, the samples were rinsed with phosphate buffer three times for 10 minutes each. Next, the fixed samples were submerged in 1% (w/v) osmium tetroxide for four hours and then washed three times with phosphate buffer for two hours each. Dehydration was performed using a graded ethanol series: 30% for one hour, 50% for one hour, 70% for one hour, 85% for two hours, 95% for two hours, and 100% for two hours, followed by overnight treatment in 100% ethanol. The samples were subsequently dried by Hitachi HCP-2 critical point dryer, coated with gold using a Hitachi E-1010 ion sputter coater, and examined using an FEI Quanta 200 scanning electron microscope at 20 KV.

### Microbialite DNA extraction, PCR, and amplicon sequence analysis

Microbialite DNA from each sample was extracted using a PowerSoil DNA isolation kit (Qiagen, Germany). The bacterial community structure was investigated by amplifying the V6-V8 region of the 16S rRNA genes using primer sets 968F (AACGCGAAGAACCTTAC) and 1391R (ACGGGCGGTGWGTRC) (23). The PCR protocol entailed 30 cycles comprising an initial step of 94℃ for 5 minutes, followed by 94℃ for 30 seconds, 52℃ for 20 seconds, 72℃ for 45 seconds, and finally 72℃ for 10 minutes. The resulting PCR products, approximately 423 bp in size, were purified using the MinElute Gel Extraction Kit (QIAGEN). Subsequently, up to 90 V6-V8 amplicon libraries were prepared and subjected to quantitative PCR (qPCR) to verify the uniformity of each library prior to pooling. Sequencing of the pooled libraries was performed on an Illumina MiSeq V2 platform using the MiSeq Reagent Kit V3 (paired end [PE]; 2 x 300 bp), conducted by Biotech Ltd., Taiwan.

Amplicon sequence analysis was performed utilizing the Quantitative Insights Into Microbial Ecology 2 (QIIME 2) software (version 2022.2) (24). Primers targeting the V6-V8 regions were removed using cutadapt (version 4.0) (25). Following primer removal, sequences underwent denoising via the DADA2 algorithm, incorporating quality filtering—sequences were trimmed to 246 base pairs for the forward reads and 259 base pairs for the reverse reads—and chimera removal (26). The qualified amplicon sequence variants (ASVs) were taxonomically classified using the classifier-consensus-vsearch plugin (27), against the SILVA 138 NR99 database (28, 29). To enhance the resolution of the ASV abundance table, ASV sequences were further clustered employing the K-mer-based taxonomic (KTU) clustering algorithm (30).

To ascertain the community profile of eukaryotes, amplicons of the V9 region of the 18S rRNA gene, approximately 315 bp in length, were generated using the primer set 1195F (AACAGGTCTGTGATG) and 1510R (CCTTCYGCAGGTTCACCTA).

The PCR conditions were consistent with those previously described. Following sequencing on the MiSeq platform, downstream bioinformatic analyses were performed similarly to those mentioned above, with the exception that length trimming was adjusted to 206 bp for forward reads and 204 bp for reverse reads.

### Statistical analysis

Alpha diversity indices, Chao1 and Shannon, were computed for all samples. Correlations between these indices and environmental factors were assessed using Pearson correlation tests. The most dominant *Cyanobacteria* family identified in the samples was subjected to phylogenetic analysis using MEGA 11 (31). Sequence alignments were performed using the ClustalW algorithm, followed by Maximum Likelihood phylogenetic inference with 500 bootstrap replicates. For microbial community interaction analysis, we utilized FastSpar v1.0 (32) to estimate correlation.

This tool quantified correlations between KTUs at the family level, determining the statistical significance of these correlations using bootstrap procedures. The resultant correlation and p-value matrices generated by FastSpar were employed to construct a network of significant correlations for each sample. Co-occurrence networks were created using the igraph (33) package in R software (34), consisting of undirected weighted networks based on statistically significant correlations (p < 0.05) greater than 0.8. These networks were visualized using Cytoscape (version 3.10.0) (35). Network analysis tools within Cytoscape calculated network properties, and networks from samples at identical depths were integrated using the ’merge network’ function. Nodes not directly connected to the dominant *Cyanobacteria* family were filtered out to refine the network structure.

## Functional pathway analysis of bacterial community

PICRUSt2 was employed to predict the potential functions of the bacterial community in each tidal pool (36). ASVs were aligned with reference 16S sequences using HMMER (37). Subsequently, EPA-NG (38) was utilized to determine the optimal placement of ASVs within the reference tree, and GAPPA (39) was used to generate a new phylogenetic tree. The new phylogenetic tree was then used to predict the copy numbers of Enzyme Classification (EC) numbers for each ASV through hidden-state prediction approaches (40). The predicted EC number abundances for each sample were calculated and normalized by read depth. MetaCyc pathway abundances were then calculated based on the EC number abundances from each sample. Ggpicrust2 (41) was applied for downstream analysis of pathway abundances. Differential abundance testing of pathways across different tidal pools was performed using the LinDA approach (42), with p-values corrected for multiple hypothesis testing using the Benjamini–Hochberg procedure (43). The top 50 pathways with significant differences between tidal pools 0 and 30 were presented in a heatmap.

## Author contribution statement

**Yunli Eric Hsieh**: Data curation, Formal analysis, Visualization, Writing – original draft. **Sung-Yin Yang**: Formal analysis, Supervision, Visualization, Writing – review and editing. **Shao-Lun Liu**: Conceptualization, Data curation, Funding acquisition, Resources, Writing – review and editing. **Shih-Wei Wang**: Conceptualization, Writing – review and editing. **Wei-Lung Wang**: Conceptualization, Writing – review and editing. **Sen-Lin Tang**: Resources, Writing – review and editing. **Shan-Hua Yang**: Conceptualization, Data curation, Funding acquisition, Investigation, Methodology, Project administration, Supervision, Visualization, Writing – original draft

## Acknowledgement

We would like to thank Prof. Po-Yu Liu and Dr. Sean Ting-Shyang Wei for instructing bioinformatic analysis. We also thank Dr. Wann-Neng Jane for providing the SEM equipment and service. We also thank Dr. Ming-Yang Ho for suggestions. This work was financially supported by National Taiwan University from Excellence Research Program - Core Consortiums (NTUCCP-113L894406) within the framework of the Higher Education Sprout Project by the Ministry of Education (MOE) in Taiwan. This study was also supported financially by the National Science and Technology Council, Taiwan (MOST 110-2611-M-002-024 and MOST 111-2611-M-002-023) to SHY, partially by the National Science and Technology Council, Taiwan (MOST 111-2621-B-029 -002 -MY3) to SLL, and partially by Kenting National Park, Taiwan to SWW. The authors declare that they have no known competing financial interests or personal relationships that could have appeared to influence the work reported in this paper.

## Data availability Statement

The sequenced data could be found at NCBI under BioProject accession number: PRJNA1164468.

